# The protein modifier SUMO is critical for Arabidopsis shoot meristem maintenance at warmer ambient temperatures

**DOI:** 10.1101/2021.02.04.429700

**Authors:** Valentin Hammoudi, Bas Beerens, Martijs J. Jonker, Tieme A. Helderman, Georgios Vlachakis, Marcel Giesbers, Mark Kwaaitaal, Harrold A. van den Burg

## Abstract

Short heat waves (>37°C) are extremely damaging to non-acclimated plants and their capacity to recover from heat stress is key for their survival. To acclimate, the HEAT SHOCK TRANSCRIPTION FACTOR A1 (HSFA1) subfamily activates a transcriptional response that resolves incurred damages. In contrast, little is known how plants acclimate to sustained non-detrimental warm periods at 27-28°C. Plants respond to this condition with a thermomorphogenesis response. In addition, HSFA1 is critical for plant survival during these warm periods. We find that SUMO, a protein modification whose conjugate levels rise sharply during acute heat stress in eukaryotes, is critical too for plant longevity during warm periods, in particular for normal shoot meristem development. The known SUMO ligases were not essential to endure these warm periods, alone or in combination. Thermo-lethality was also not seen when plants lacked certain SUMO proteases or when SUMO chain formation was blocked. The SUMO-dependent thermo-resilience was as well independent of the autoimmune phenotype of the SUMO mutants. As acquired thermotolerance was normal in the *sumo1/2* knockdown mutant, our data thus reveal a role for SUMO in heat acclimation that differs from HSFA1 and SIZ1. We conclude that SUMO is critical for shoot meristem integrity during warm periods.

**Highlight:** The protein modifier SUMO governs shoot meristem maintenance in Arabidopsis allowing sustained rosette development when plants endure a sustained warm non-detrimental period of 28 degrees Celsius.

## Introduction

Plants are constantly challenged by temperature fluctuations caused by day-night cycles, weather conditions, seasonal shifts, climate extremes and global warming. These fluctuations differ in temperature range, duration and gradient. Plants acclimate to the ambient temperature via an intertwined molecular network that resolves the (protein) damage incurred while changing their morphology and performance to better deal with temperature extremes (Dickinson *et al.*, 2018; Ding *et al.*, 2020). Heat extremes cause among others protein unfolding, membrane damage, and release of reactive oxygen species (ROS) (Hightower, 1991). Heat-inflicted damage is perceived via Heat Shock transcription Factors (HSFs) that up-regulate expression of Heat Shock Proteins (HSPs) and other protein-folding chaperones (Wu, 1995). The misfolded proteins are then stabilized and refolding by HSPs and, when ineffective, HSPs target misfolded proteins for degradation (Wang *et al.*, 2004). In particular, HSP70 and HSP90 are central players for resolving protein damage, while HSP101 is critical for short- and long-term acquired thermotolerance (SAT and LAT) (Hong and Vierling, 2001; Queitsch *et al.*, 2000).

In Arabidopsis (*Arabidopsis thaliana*) heat stress is perceived by the four members of the HSF clade A1 (HSFA1) (Liu and Charng, 2013; Liu *et al.*, 2011; Ohama *et al.*, 2016). Prior to heat stress, HSFA1 is kept in an inactive state in the cytoplasm by HSP70 and HSP90 (Ohama *et al.*, 2016; Yoshida *et al.*, 2011). Upon heat stress, HSFA1 shuttles to the nucleus where it induces expression of heat-responsive genes including genes encoding additional transcription factors (*HSFA2, DREB2A* (*DEHYDRATION RESPONSIVE ELEMENT BINDING 2A*), *DREB2B*, and *MBF1C* (*MULTIPROTEIN BRIDGING FACTOR 1C*). Combined these gene products form a second transcriptional wave that orchestrates the stress response while promoting plant acclimation to a subsequent heat wave (Kotak *et al.*, 2007; Liu *et al.*, 2011; Nishizawa-Yokoi *et al.*, 2011; Yoshida *et al.*, 2011). Overall, HSFA1 is regarded to be the master regulator of heat stress in plants. HSFA1 plays also a role in cold acclimation via NPR1 (NON-EXPRESSER OF PATHOGENESIS-RELATED GENES 1), which is a master regulator of the plant response to biotrophic pathogens (Olate *et al.*, 2018).

Importantly, protein modifications play as well an important regulatory role in response to heat stress, in particular sumoylation. Many proteins are for example SUMO (Small ubiquitin-like modifier) modified when eukaryotic cells experience acute heat stress (Golebiowski *et al.*, 2009; Miller *et al.*, 2010; Miller *et al.*, 2013; Miller and Vierstra, 2011; Tatham *et al.*, 2011). One mechanism is that sumoylation promotes the solubility of proteins during heat stress in mammalian cells (Liebelt *et al.*, 2019). In human cells sumoylation also controls the transcriptional response to heat stress by modifying the transcription factor HSF1, the closest homolog of Arabidopsis HSFA1 (Hietakangas *et al.*, 2003; Hilgarth *et al.*, 2003; Hong *et al.*, 2001). Also for Arabidopsis evidence exists that the HSF regulatory pathway is subject to sumoylation. For instance, the transcription factors HSFA1b, HSFA1d, HSFA2 and DREB2A are sumoylated *in planta* (Augustine and Vierstra, 2018; Miller et al., 2010; Miller et al., 2013; Rytz et al., 2018), but the role of SUMO in *(a)* acute heat stress and *(b)* acclimation to sustained warm periods remains poorly understood in plants. SUMO conjugate levels increase sharply in Arabidopsis in response to heat stress affecting hundreds of targets (Miller *et al.*, 2013; Rytz *et al.*, 2018). This global increase depends strongly on the SUMO E3 ligase SIZ1 (SAP AND MIZ-FINGER DOMAIN 1) and is attenuated in plants overexpressing HSP70 (Kurepa *et al.*, 2003; Miller *et al.*, 2013; Rytz *et al.*, 2018; Yoo *et al.*, 2006). The latter implies that heat stress triggers a global increase in SUMO conjugate levels that is connected to protein damage via an unknown mechanism and with an unknown impact on plant development and stress acclimatization. Studies with mammalian cells have suggested that acute heat stress causes inactivation of certain SUMO proteases, which would explain the sudden increase in SUMO adducts (Liebelt *et al.*, 2019; Pinto *et al.*, 2012).

In contrast to heat stress, sustained warm periods of 27-28°C do not lead to permanent protein damage in plants and they do not cause up-regulation of known heat stress marker genes (Kumar and Wigge, 2010). Instead, plants respond to warm temperatures by altering their development (called thermomorphogenesis), including early flowering, leaf hyponasty, hypocotyl stretching and petiole elongation. (Casal and Balasubramanian, 2019; Qiu *et al.*, 2019; Quint *et al.*, 2016). As a consequence, the rosette of Arabidopsis adopts an open architecture, which is supposed to increase the leaf cooling capacity. This thermomorphogenesis response depends as well on SIZ1 activity and the two main SUMO isoforms, SUMO1 and −2 (Hammoudi *et al.*, 2018).

All our studies with the SUMO1/2 knockdown mutant (*sumo1-1;35S*_*Pro*_*::amiR-SUMO2*, hereafter *sumo1/2*^*KD*^) have thus far indicated that it closely resembles the phenotype of *siz1* loss-of-function mutants, including *(i)* high levels of the defense hormone salicylic acid (SA), *(ii)* constitutive expression of PATHOGENESIS-RELATED PROTEINS 1 and −2 (PR1 and PR2), *(iii)* spontaneous cell death, *(iv)* dwarf stature with curled leaves, *(v)* early flowering, *(vi)* loss of apical dominance, but also *(vii)* compromised thermo- and photomorphogenesis responses (Hammoudi *et al.*, 2018; van den Burg *et al.*, 2010). The autoimmune phenotype of *siz1* is dependent on the immune receptor SNC1 (SUPPRESSOR OF *NPR1-1*, CONSTITUTIVE 1) (Gou *et al.*, 2017). Normally, SNC1 autoimmunity is suppressed at 28°C, resulting in a wild type rosette shape at this temperature (Yang and Hua, 2004; Zhu *et al.*, 2010). Autoimmunity due to the *siz1* mutation is, however, not suppressed at 28°C and correspondingly growth of *siz1* rosettes is only partially recovered at 28°C (Hammoudi *et al.*, 2018). This signifies that SIZ1-dependent SUMO conjugation inhibits SNC1 immune signaling, directly or indirectly, both at normal and warm temperatures.

We now report that SUMO1/2 combined are critical for sustained rosette development when plants endure periods of 28°C. Interestingly, this role of SUMO1/2 is highly reminiscent of the function of Arabidopsis HSFA1 at this temperature (Liu and Charng, 2013; Liu *et al.*, 2011). Yet the HSFA1 response appears to be intact the *sumo1/2*^*KD*^ mutant. Moreover, the SUMO protein levels are critical for rosette development and shoot meristem integrity at 28°C independent of SIZ1 and HPY2, the two main SUMO E3 ligases in Arabidopsis. This role of SUMO in thermo-resilience did also not depend on immune signaling. We thus report a novel role for SUMO in plant thermo-resilience.

## Materials and methods

### Plant materials and growth conditions

*Arabidopsis thaliana* (L.) Heynh was used for the experimentation with the mutants and transgenic lines as detailed in the **Supplementary Table S1**. All lines were obtained from Nottingham Arabidopsis Stock Centre (NASC) or from the sources listed in the **Supplementary Table S1**. To obtain additional *sumo1-1;35S*_*Pro*_*::amiR-SUMO2* lines (*sumo1/2*^*KD*^), independent *35S*_*Pro*_*::amiR-SUMO2* (T1) lines were screened for low *SUMO2* expression using real time RT-PCR. Four additional lines with low *SUMO2* expression levels were identified and they were crossed with *sumo1-1;sumo3-1.* After crossing of *sumo1/2*^*KD*^ line B with *pad4-1, sid2-1* or *eds1-2* with, the F2 progeny was genotyped for the alleles *SUMO1, sumo1-1*, *amiR-SUMO2* in *PFK7* and *amiR-SUMO2* in *CYP98A3*_*pro*_ alleles according to (Hammoudi *et al.*, 2017). Primers and genotyping details are given in **Supplementary Table S2 and S3**, respectively.

Arabidopsis plants were grown under short day conditions (11 h light/13 h dark) at a constant temperature of 22°C or 28°C unless specified otherwise. When grown on plates, seeds were gas-sterilised, stratified in liquid for 2 or 3 days at 4 °C in the dark conditions and then germinated on 0.5x Murashige and Skoog salt mixture with Gamborg B5 vitamins (Duchefa), 0.5 g MES (Duchefa), 0.1 g Myo-Inositol (Merck) and 0.8 or 1% Daishin agar (Duchefa) pH 5.7, with the same light regime at 22 °C. To induce proteotoxic or abiotic stress, the plates were supplemented with L-Canavanine (10 μM), mannitol (300 mM), or NaCl (75 mM). Heat stress treatments were performed as previously described (Liu and Charng, 2013). Briefly, for short-term acquired thermotolerance (SAT) 7-day-old seedlings were acclimated at 37°C for 60 min and then allowed to recover for 120 min at 22°C before being incubated at a noxious temperature of 44°C for 90 min. For long-term acquired thermotolerance (LAT), 7-day-old seedlings were acclimated at 37°C for 60 min, then recovered for two days at 22°C before being incubated at 44°C for 50 min.

### Gene expression quantification

For the gene expression analysis, total RNA was isolated using TRIzol Reagent (ThermoFisher) according to the suppliers’ instructions. A total of 2 μg RNA was used for cDNA synthesis using Superscript III (ThermoFisher). RNAse activity was inhibited by adding RiboLock (ThermoFisher) during cDNA synthesis. Real-time PCR was performed on an ABI 7500 real-time PCR system (Applied Biosystems). Primers used for the gene expression analysis are given in **Supplementary Table S2.** The cycling program was set to 2 min, 50°C; 10 min, 95°C; 40 cycles of 15 s at 95°C; and 1 min, 60°C, and a melting curve analysis was performed at the end of the PCR. All primer pairs were tested for specificity and for amplification efficiency using a two-fold dilution series of a mixed cDNA sample. The biological samples were normalized against three housekeeping genes (Czechowski *et al.*, 2005) using geometric averaging: *ACT2* (At3g18780), *UBQ10* (At4g05320), and *UBC21* (At5g25760). Primers used were described previously (Czechowski *et al.*, 2005) (**Supplementary Table S2**). The data were analysed using the methods included in the gene expression software qBASE+ (BioGazelle, Belgium) with a correction for the amplification efficiencies of the primer pairs.

### Protein analysis

For the heat shock treatments, Arabidopsis seedlings were pre-grown on plates (half-strength Murashige and Skoog salt mixture supplemented with Gamborg B5 vitamins) for 14 days under SD light conditions at 22°C. Seedlings were exposed to a 30-minute acute heat shock at 37°C in the dark and left to recover at 22°C for 60 min before freezing the samples in liquid nitrogen for storage at −80°C untill total protein extraction (Kurepa *et al.*, 2003). The total protein fraction was extracted by homogenizing the frozen plant material with metal beads (2.5 mm diameter) in a TissueLyser II (Qiagen). Per fresh sample weight, two volumes of a freshly prepared Protein extraction (PE) buffer (50 mM potassium phosphate buffer pH 7.0, 150 mM NaCl, 1 mM EDTA, 2% (w/v) poly(vinylpolypyrrolidone), 1× cOmplete mini EDTA-free protease inhibitor cocktail (Roche), 1% Nonidet P-40, 10% (w/v) glycerol and either 5 mM DTT or 20 mM NEM) were added followed by mixing with a vortex for 10 sec. The cell lysates were incubated for 15-30 minutes (4°C, 20 rpm on rotating wheel) and then centrifugated for 10 min at 16,000 *g* (4°C). Total protein concentrations were measured using a BCA protein assay kit (Sigma) and normalized when needed by adding extra PE buffer. For denaturation, the protein extracts were mixed 1:1 with 2× Sample loading buffer (100 mM Tris-HCl pH 6.8, 4% SDS, 20% glycerol, and 100 mM DTT) and then boiled for 10 min. The denatured protein samples (20 μL) were loaded on a 10-15% SDS-PAGE gels, separated by electrophoresis, and then transferred onto a polyvinylidene fluoride membrane (PVDF; Immobilon-P, Millipore) using semi-dry blotting according to the suppliers’ instructions (Hoefer). Equal protein loading and transfer to the blot was confirmed by staining the PVDF membranes with 0.1% Ponceau S in 5%(v/v) acetic acid. After destaining with two rinses of 5% (v/v) acetic acid, the blot was scanned and further destained by rinsing them thrice with Tris Buffer Saline (TBS; 25 mM Tris-HCl pH 7.6, 137 mM NaCl, and 2.7 mM KCl). The membranes were blocked with 5% (w/v) skimmed milk powder dissolved in TBS supplemented with 0.05% v/v Tween 20 (TBST). Incubations with both the primary and secondary antibodies were performed in TBST supplemented with 5% milk powder, followed by three rinses of 5, 10 and 15 minutes with TBST (after the primary antibody) or TBS (after the secondary antibody). The secondary immunoglobulins conjugated to horseradish peroxidase were visualized using enhanced chemiluminescence with a commercial kit (ECL Pierce Plus, ThermoFisher) or a homemade solution (100 mM Tris-HCl pH 8.5, freshly supplemented with 50 μL of 250 mM Luminol in DMSO, 22 μl of 90 mM coumaric acid in DMSO, and 3 μl 30% H_2_O_2_ solution). The luminescence signals were detected using MXBE Kodak films (Carestream). Details on the antibodies used are given in the **Supplementary Table S1**.

### Microarray gene expression analysis

Briefly, *pad4-1*, *sumo1/2*^*KD*^*;pad4-1*, *siz1-2;pad4-1* and *eTK* plants were grown on soil at 22°C in short day conditions for 2 weeks and then transferred to 28°C. Leaf samples were collected at moment of the shift to 28°C and after 24 or 96 h at 28°C and used for RNA extraction using Trizol Reagent (ThermoFisher). Three biological replicates were collected for each sample. Total RNA was further purified using the RNAeasy Plant mini Kit (Qiagen). The RNA quality was examined by monitoring the Absorbance ratios at 260/280 nm and 260/230 nm. A total of 100 ng total RNA was amplified using the GeneChip WT PLUS Reagent Kit (Affymetrix) to generate biotinylated sense-strand DNA targets. The labeled samples were hybridized to Affymetrix Arabidopsis Gene 1.1 ST arrays (ThermoFisher). Washing, staining and scanning was performed using the GeneTitan Hybridization, Wash, and Stain kit for WT Array Plates, and the GeneTitan instrument (Affymetrix). All arrays were subjected to a set of quality control checks, such as visual inspection of the scans, checking for spatial effects through pseudo-color plots, and inspection of pre- and post-normalized data with box plots, ratio-intensity plots and principal component analysis (PCA). Normalized expression values were calculated using the robust multi-array average (RMA) algorithm (Irizarry *et al.*, 2003). The experimental groups were contrasted to test for differential gene expression. Statistical analysis was performed to test the experimental groups for differential gene expression compared to *pad4-1*, at each time point. Empirical Bayes test statistics were used for hypothesis testing (Smyth, 2004) using the Limma package in R 3.4.1 (http://cran.r-project.org/), and all obtained *p*-values were corrected for false discoveries according to Storey and Tibshirani (Storey and Tibshirani, 2003). The DEGs were selected using an F-test in the Limma package, designed to test for differential expression between any of the four genotypes (*pad4-1, pad4-1;sumo1/2*^*KD*^*; pad4-1;siz1-2; pad4-1* and *eTK* plants) at the individual time points. An overall comparison of the gene expression responses due to increase in ambient temperature between all genotypes simultaneously was made by PCA. To interpret the PCA results, a gene ontology (GO) analysis (on “biological processes” only) was performed on the top 500 genes with the highest absolute loading scores for PC1 and PC2, respectively, using AgriGOv2 (Tian *et al.*, 2017). This PCA+GO comparison was also made based on the genome wide response of (a) all measured genes and also based on the transcriptome response of a collection of genes, selected using an F-test in the Limma package, designed to test for differential expression between any of the four groups. The AgriGOv2 results were plotted in R using clusterProfiler (Yu *et al.*, 2012). The p-values for the GO terms were calculated using a hypergeometry test with Yekutieli FDR under dependency; q-value (P_adj_) <0.01 using the complete GO as implement in AgriGOv2. The dot plots displaying the significant GO terms for the different PC axes were generated using the R package clusterProfiler (Yu *et al.*, 2012). Absence of a dot signifies that GO terms is non-significant at that time point.

### Cryo-scanning electron microscopy (cryo-SEM)

Excised apical meristems were attached to a sample holder using a very thin layer of Tissue-Tek compound (EMS, Washington, PA, USA). Samples were plunge frozen in liquid nitrogen and subsequently placed in a cryo-preparation chamber (MED 020/VCT 100, Leica, Vienna, Austria). To sublimate any water vapor contamination (ice) from the surface, the samples were kept for 4 minutes at −93°C. Samples were then sputter coated with a 12 nm layer of Tungsten (W), and transferred under vacuum to the field emission scanning electron microscope (Magellan 400, FEI, Eindhoven, the Netherlands) onto the sample stage at −120°C. The images were taken with SE detection at 2 kV, 13 pA.

### Propidium iodide staining and imaging of roots

Seeds were germinated for 4 days on vertical 0.5x MS plates at 22 °C. Seedlings that germinated were transferred to new 0.5x MS plates and grown for another 8-10 days at *(i)* 22°C, *(ii)* 28°C, or *(iii)* first placed at 22°C for 3 days and then shifted to 28°C for another 5-7 days. These plates were scanned at 12 days post germination and the main root length and number of lateral roots was measured in ImageJ (https://fiji.sc/) for each sample. For confocal laser scanning microscopy (CLSM), the roots were detached from the seedlings with a scalpel and transferred to a Propidium iodide (PI) (ThermoFisher) staining solution (final concentration 10 μg/ml, diluted from a 1 mg/ml stock) in a 6-well plate. Roots were incubated for 5-10 min in the dark, rinsed in a separate well with water, transferred to a microscopy slide and covered with a coverslip. Imaging was performed using a Zeiss LSM 510 or Nikon A1 CLSM, using a 20x Plan-Apochromat lens with Numerical Aperture (NA) 0.75 or a 20x Plan Fluor DIC N2 lens with an NA of 0.75, respectively. PI was excited with a 561nm diode laser, thereafter the emission light was selected using a 600/30 nm or 595/50 nm bandpass filter, respectively, before detection with photomultiplier tubes. Simultaneously, a bright field image was recorded with the transmitted light detector.

### Quantification and Statistical analysis

Data visualization and statistical analyses of the data were performed using the software PRISM (GraphPad). Statistical analyses and plots were computed in PRISM using built-in functions. Statistical information is further specified in the Figure captions.

## Results

Previous work on *sumo1/2*^*KD*^ exposed that its phenotype resembles the SUMO ligase mutant *siz1-2* (Hammoudi *et al.*, 2018; van den Burg *et al.*, 2010). While studying the role of SUMO in thermomorphogenesis (**Fig. 1A**), we noted that development of *sumo1/2*^*KD*^ seedlings was arrested at 28°C resulting in their collapse 14 days post germination. Opening of the cotyledons was still normal at this temperature, but *sumo1/2*^*KD*^ failed to develop new leaves. In contrast, at normal temperatures (22°C) new leaves formed, but they were curled and small. At 16°C, leaf development of *sumo1/2*^*KD*^ was similar to 22°C except that the rosette diameter doubled in size. A similar cool-temperature response (*i.e*. increased rosette diameter at 16°C) was seen for two SNC1-dependent autoimmune mutants, *srfr1-4* (*suppressor of rps4-rld* 4) and *bon1* (*bonzai 1*). This latter observation suggests that the cool temperature response might be connected with unbalanced SNC1 signaling, while the high temperature collapse clearly differs from the SNC1-dependent phenotype of *siz1*. At 25°C (**Fig. 1A**), *sumo1/2*^*KD*^ seedlings survived even though their growth was suppressed compared to 22°C.

**Fig. 1.**
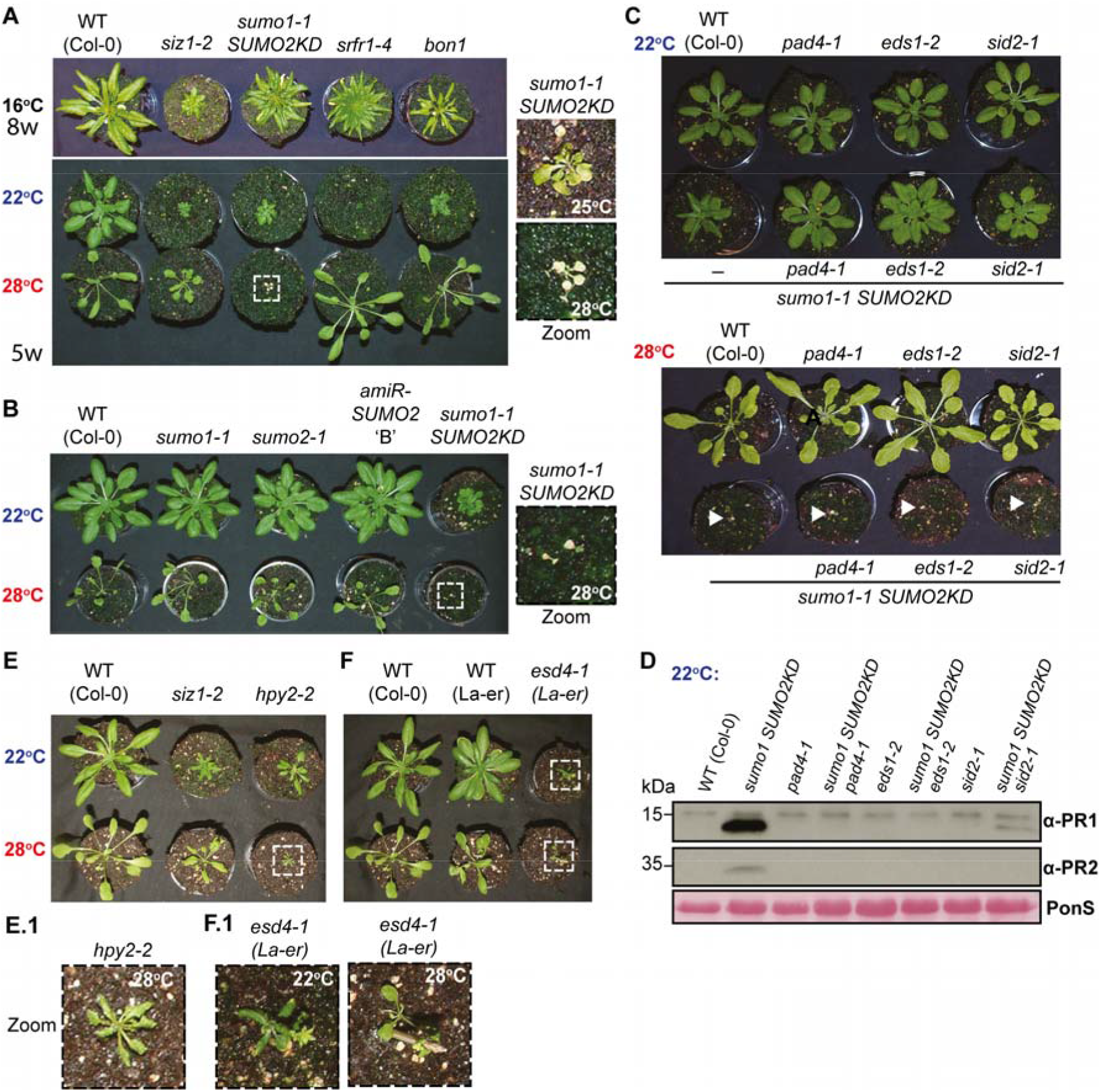
SUMO and SUMO2 combined are essential for Arabidopsis to sustain elevated temperatures. **A.** Growth phenotype of the indicated plant genotypes at different temperatures to assess suppression of SNC1-dependent autoimmunity at 28°C. *bon1* and *srfr1-4* are two mutants with SNC1-dependent autoimmunity. Plant age is indicated on the left (w, weeks). Images on the right show the rosette phenotype of *sumo1-1;SUMO2*^*KD*^ at 25 vs. 28°C. WT, wildtype accession Col-0. **B.** Growth phenotype at 22/28°C for the single mutants *sumo1-1*, *sumo2-1* and *SUMO2*^*KD*^ (line B) and the corresponding double mutant *sumo1-1;SUMO2*^*KD*^. Picture was taken four weeks after germination. **C.** Premature collapse of *sumo1-1;SUMO2*^*KD*^ at 28°C (bottom) is independent of EDS1, PAD4, or SA accumulation. At 22°C (top), the rosette morphology *sumo1-1;SUMO2*^*KD*^ was partially recovered in the *eds1-2, pad4-2, sid2-1* backgrounds. Picture was taken five weeks post germination. **D.** Immunoblot showing PR1 and PR2 protein levels in 5-week-old plants. PR1 and PR2 levels were suppressed when *sumo1-1;SUMO2*^*KD*^ is introduced in the *pad4-1*, *eds1-2* or *sid2-1* backgrounds. **E.** Null mutant of the SUMO E3 ligases SIZ1 and HPY2 survive at 28°C (**E.1**, zoom of *hpy2-2* rosette). **F.** Null mutant of the SUMO protease ESD4 (*esd4-1*) survives at 28°C (**F.1**, zoom of *esd4-1* rosette).

To exclude that SUMO-dependent thermo-resilience is an artefact of the plant transformation procedure, we assessed thermo-resilience of the mutants *sumo1-1*, *sumo2-1,* and *35S*_*Pro*_*::amiR-SUMO2* (line B). Each mutant withstood 28°C without tissue collapse while also displaying a wildtype-like thermomorphogenesis response as evidenced by the elongated petioles and leaf blades (**Fig. 1B**). Moreover, we generated four independent crosses between *sumo1-1* and plants with reduced *SUMO2* transcript levels due to expression of *amiR-SUMO2* (lines P, S, V, and W). None of the *amiR-SUMO2* parental lines had an aberrant growth phenotype at 22°C, while plants with a typical *sumo1/2*^*KD*^ morphology were present in the F2 progeny*, i.e.* plants with a dwarf stature, curled leaves, lesions and premature flowering (**Supplementary Fig. S1A**)(van den Burg *et al.*, 2010). Genotyping of the dwarf plants confirmed they were homozygous for *sumo1-1* and they contained (at least) one gene copy of *35S*_*Pro*_*::amiR-SUMO2*. Similar to the original *sumo1/2*^*KD*^ mutant, the dwarf plants were thermo-sensitive resulting in their collapse after two weeks at 28°C. Clearly, SUMO1/2 combined confer thermo-resilience at 28°C.

Since the developmental phenotype of *siz1* and *sumo1/2*^*KD*^ is in part caused by autoimmunity and concomitantly high SA levels (Hammoudi *et al.*, 2018; Lee *et al.*, 2007; van den Burg *et al.*, 2010), *sumo1/2*^*KD*^ was crossed with mutants defective in this immune response, *i.e. pad4-1* (*phytoalexin-deficient4-1*), *eds1-2* (*enhanced disease susceptibility1-2*) and *sid2-1* (*salicylic acid induction deficient2-1*). The proteins PAD4 and EDS1 form together an immune hub, while *SID2* encodes for the enzyme ISOCHORISMATE SYNTHASE 1 (ICS1) needed for SA biosynthesis in response to pathogen recognition. As expected, *sumo1/2*^*KD*^ autoimmunity was suppressed at 22°C in the backgrounds *pad4*, *eds1* and *sid2* (**Fig. 1C**). For example, the levels of the defense marker proteins PR1 and PR2 were no longer elevated in the triple mutants *sumo1/2*^*KD*^*;pad4, sumo1/2*^*KD*^*;eds1* and *sumo1/2*^*KD*^*;sid2* (**Fig. 1C, 1D**). Nonetheless, these triple mutants still collapsed when grown at 28°C, while the single mutants *pad4, eds1* and *sid2* developed normally at 28°C (including a normal thermomorphogenesis response). Thus, thermo-lethality of *sumo1/2*^*KD*^ is independent of its autoimmunity.

To assess whether other components of the SUMO (de)conjugation pathway support sustained plant growth at 28°C, different loss-of-function mutants were tested including the isoform SUMO3, the SUMO E3 ligase HPY2/MMS21 (HIGH PLOIDY 2, MMS21) and different SUMO proteases. As SUMO proteases, the role of ESD4 (EARLY IN SHORT DAYS 4) alone, OTS1/2 (OVERLY TOLERANT TO SALT 1 and 2) combined, and SPF1/2 (SUMO PROTEASE RELATED TO FERTILITY 1 and 2) combined were tested (**Fig. 1E, 1F, Supplementary Table S3**). None of the corresponding proteins appeared to be essential for survival at 28°C. Likewise, SUMO E4 ligase activity mediated by PIAL1/2 (PROTEIN INHIBITOR OF ACTIVATED STAT LIKE 1 and 2) was not required for growth at 28°C, alone or in combination with SIZ1 (using the triple mutant *siz1;pial1;pial2*) (**Supplementary Fig. S1B**). Also three complementation lines were examined that express a variant of the E2 SUMO conjugating enzyme SCE1, SCE1(K15R), from its endogenous promoter in the *sce1-5* loss-of-function mutant background (*SCE1*_*Pro*_*::SCE1(K15R)*;*sce1-5*) (Tomanov *et al.*, 2018). These lines no longer form poly-SUMO chains, but SUMO is still attached as monomer onto acceptor lysines. Loss of SUMO chain formation did not result in thermo-lethality (**Supplementary Fig. S1C**). Thus, none of the other mutants in the SUMO machinery displayed thermo-lethality. Except for the mutants with dwarf rosettes (*siz1-2, hpy1* and *eds4*), the SUMO machinery mutants tested displayed all a normal thermomorphogenesis responses. This signifies that SUMO1/2 protein itself or mono-sumoylation of one or more substrates permits Arabidopsis to survive 28°C independent of SIZ1.

### Seven-day period at 28°C triggers thermo-lethality in *sumo1*/*2*^*KD*^ and the *hsfA1* mutant *eTK*

To assess if thermo-lethality is connected to protein damage, we determined whether Arabidopsis mutants of known damage response regulators confer thermo-resilience (**Supplementary Table S3**). Only one of these mutants showed thermo-lethality at 28°C, namely the mutant *eTK* that lacks three of the HSFA1 isoforms (*hsfA1a,b,d*) (**Fig. 2**). *eTK* was already known to collapse at 28°C (Liu and Charng, 2013). The remaining family member, HSFA1e, can apparently not compensate for thermo-sensitivity at 28°C and this isoform contributes also the least to thermotolerance (Liu and Charng, 2013; Liu *et al.*, 2011). In contrast, the mutant *hsfA2* developed normally at 28°C (**Supplementary Fig. S1D, Supplementary Table S3**). This is remarkable as the *HSFA2* gene is a major transcriptional target of HSFA1 in response to heat stress (Liu and Charng, 2013).

**Fig. 2.**
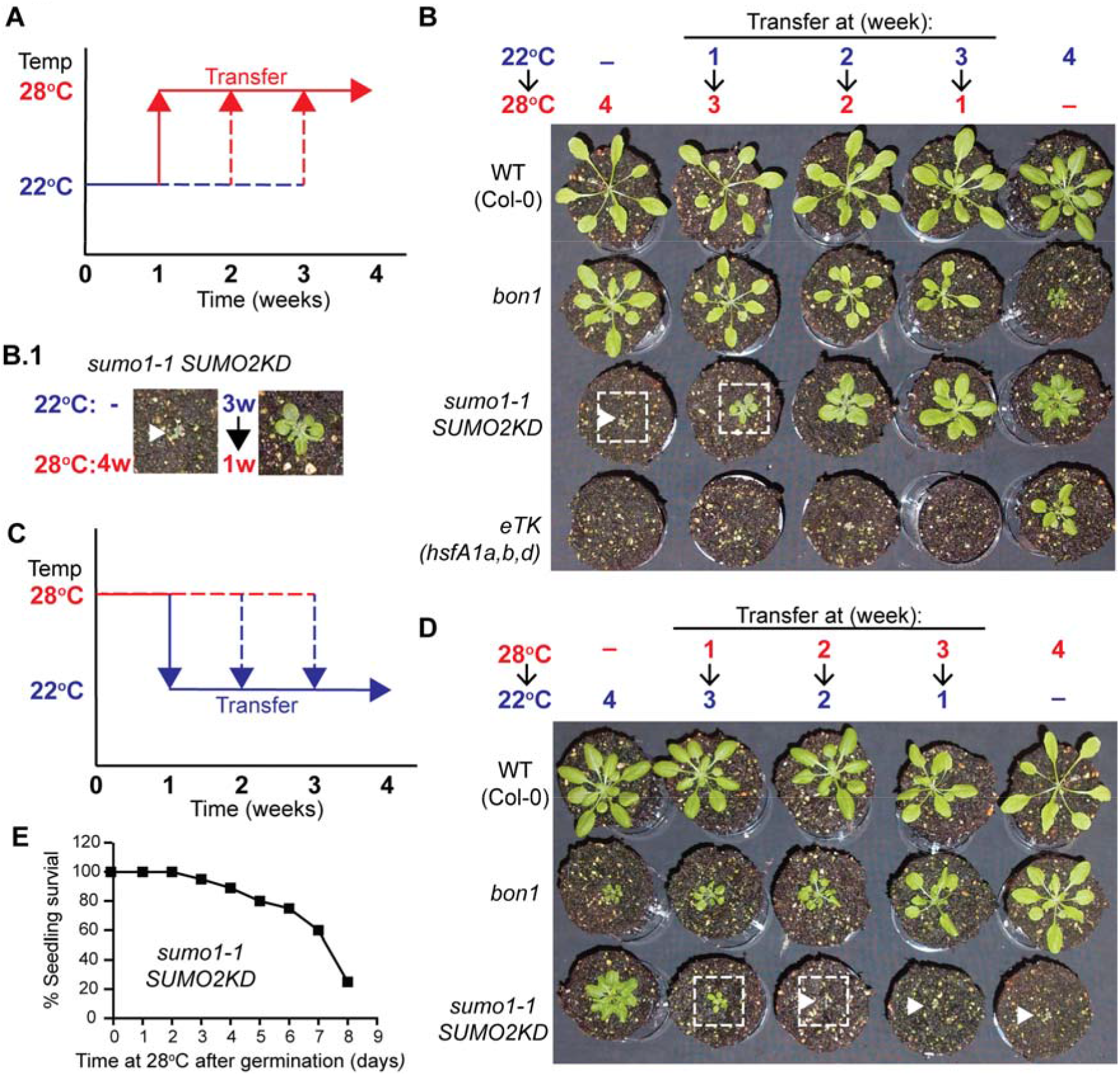
An incubation period of at least one week at 28°C results in thermo-lethality of *sumo1-1;SUMO2*^*KD*^. **A.** Diagram depicting the experimental procedure shown in (B). Following germination at 22°C, plants were transferred to 28°C after 1, 2, or 3 weeks. Control plants remained at 22 or 28°C constant temperature for four weeks. **B.** Growth of *sumo1-1;SUMO2*^*KD*^ for 1 week at 22°C is enough to prevent collapse during an additional 3 weeks at 28°C (**B.1**, zoom), while *eTK* still collapsed even when it only experienced the final week at 28°C. *bon1* was included as a control for the temperature-sensitive recovery of the growth phenotype. Picture was taken four weeks post germination (n=8 plants per line per treatment). **C.** Similar to (A), except that plants were germinated at 28°C and then transferred to 22°C. **D.** The same experiment as (B), except that it started at 28°C. The mutant *sumo1-1*;*SUMO2*^*KD*^ survived its first week at 28°C (n=3 of 24 plants), while *eTK* does not germinate at 28°C (see also Fig. 6A). *bon1* showed progressive recovery with increasing time spend at 28°C. **E.** Bar graph showing the proportion of surviving seedlings (%) of *sumo1-1;SUMO2*^*KD*^ in response to an initial growth phase at 28°C (days) followed by a shift to 22°C at the indicated day. The plants were scored after four weeks (n=28).

We compared thermo-sensitivity of *sumo1/2*^*KD*^ and *eTK* by varying the length of the warm period at different developmental stages. First, the mutants were germinated at a normal (22°C) or warm temperatures (28°C) and then after 7, 14 or 21 days the plants were shifted to the opposite temperature and their survival was scored at 28 days (**Fig. 2A, 2C**). As positive control for a ‘temperature-sensitive growth phenotype’, the mutant *bon1* was included as its autoimmune dwarf phenotype fully recovers at 28°C (Yang and Hua, 2004). Indeed, the size of *bon1* increased with more time at 28°C (**Fig. 2B**). Strikingly, an initial *‘*cool’ period of one week at 22°C was sufficient to prevent thermo-lethality of *sumo1/2*^*KD*^ even when this was followed by three weeks at 28°C although the rosette remained compact (**Fig. 2B, 2B.1**). When *sumo1/2*^*KD*^ was first grown at 22°C for two weeks followed by a warm period of two weeks, its rosette adopted a flat morphology while the rosette size increased. This change in morphology suggested that auto-immunity was partially suppressed by the warm temperature. These *sumo1/2*^*KD*^ plants still failed to show petiole and hypocotyl elongation in response to 28°C, which confirms again that the thermomorphogenesis response is compromised (Hammoudi *et al.*, 2018). In contrast to *sumo1/2*^*KD*^, *eTK* seedlings already collapsed when they experienced only 28°C during the final week of the experiment (week 4)(**Supplementary Fig. S1E**). Apparently, thermo-lethality of *eTK* is independent of its developmental stage, *i.e.* it occurred upon germination and when the plants were three weeks-old.

In the reverse experiment, *i.e.* warm start followed by a cool period (**Fig. 2C and D**), recovery of the *bon1* dwarf phenotype was again more pronounced when the warm period lasted longer. In contrast, *sumo1/2*^*KD*^ collapsed as soon as the warm period lasted two weeks or more. However, several *sumo1/2*^*KD*^ plants survived when the warm start was only one week (3/24 survivors). The *eTK* seedlings died already when they experienced one week at 28°C. We then quantified the survival rate of the seedlings in response to a warm start that lasted one to maximum eight days, before returning them to a normal temperature regime of 22°C (**Fig. 2E**). The survival rate was still >75% when *sumo1/2*^*KD*^ experienced a period of maximum five days at 28°C. After seven or eight days at 28°C, the survival rate had dropped to 60% and 25%, respectively, suggesting that the median lethal dose (LD_50_) of a warm period was approximately seven days for *sumo1/2*^*KD*^.

### A warm pulse of seven days reflects a lethal dose for *sumo1*/*2*^*KD*^ and *eTK*

To evaluate whether growth of *sumo1/2*^*KD*^ and *eTK* resumes after a warm period, seeds were germinated at a normal temperature (22°C). After two weeks, the seedings received a warm period (28°C) of variable length (2 to 14 days) after which they returned to 22°C for the remainder of the experiment (**Fig. 3A-C**). In parallel, control plants were placed at a constant temperature (22°C or 28°C), and one set remained at 28°C upon the shift to this temperature (22°C→28°C). As internal control for the temperature treatments, *bon1* and *siz1* were included again (**Fig. 3D**). As expected, *bon1* and *siz1* showed progressively more growth recovery with more time spent at 28°C (**Fig. 3D**), confirming that the temperature treatments were effective.

**Fig. 3.**
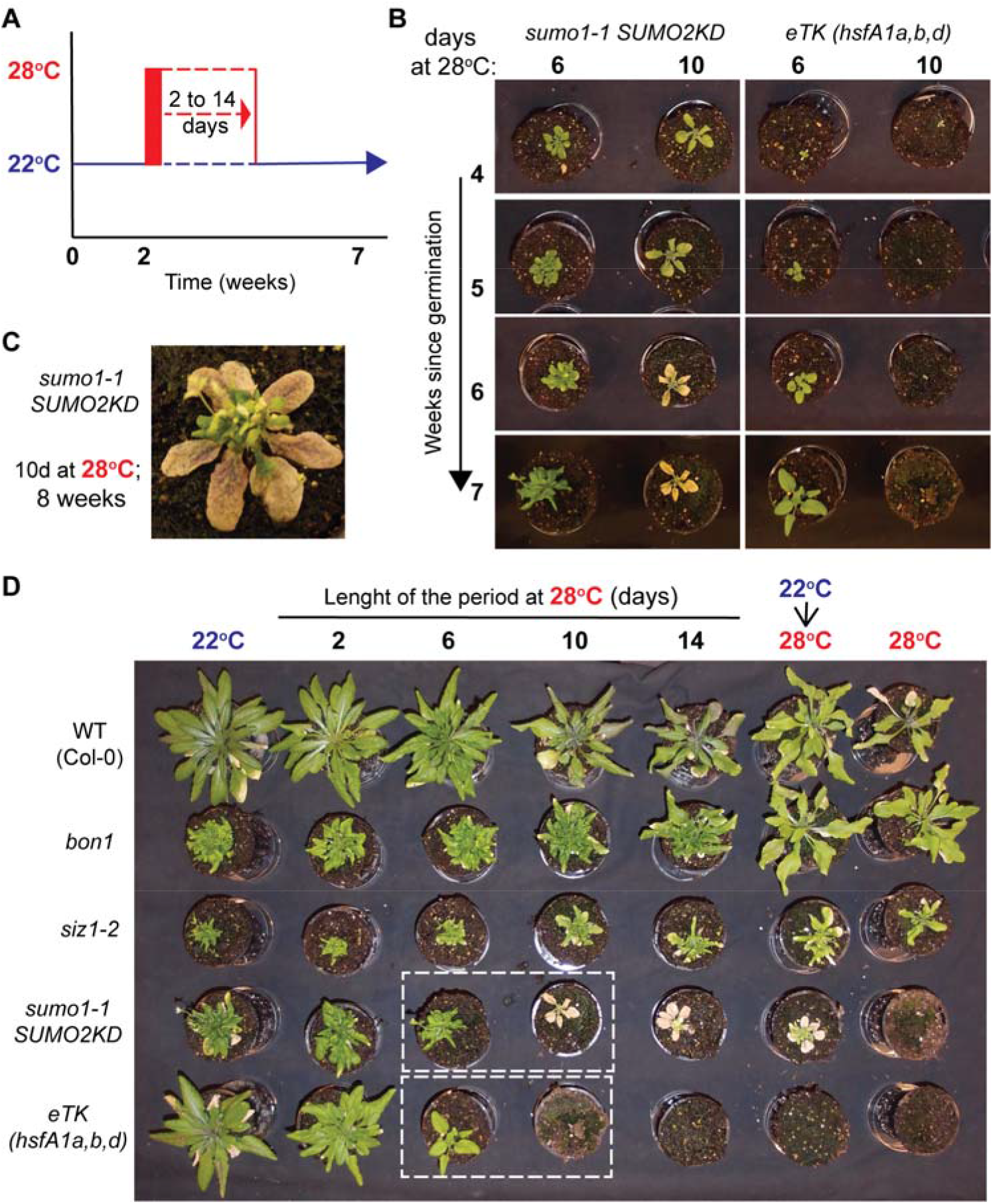
One week at 28°C results in sustained arrested development of *sumo1-1;SUMO2*^*KD*^ culminating in rosette senescence and lethality. **A.** Diagram depicting the experimental procedure. Two weeks post germination eight plants per genotype were shifted to 28°C for a period of 2-14 days, after which they received a cooler temperature regime (22°C) for another 4-6 weeks. Their development was weekly assessed starting when they were four-weeks old. **B.** Rosette development of *sumo1-1;SUMO2*^*KD*^ was arrested while *eTK* rapidly collapsed when these genotypes received a ten-day period at 28°C, but not for a six-day period. The pictures show the same plants in time (weeks) after they had experienced a brief warm period of six or ten days. **C.** Zoom of 8-weeks-old *sumo1-1;SUMO2*^*KD*^ plant that received 10 days at 28°C. The rosette stopped developing and the leaves turn eventually necrotic, while the shoot apical meristem develops a tiny, distorted inflorescence with maximum four flowers. **D.** Image depicting 7-weeks-old plants for five genotypes (left) after receiving different temperature regimes (top). Growth of both *bon1* and *siz1-2* recovered partially in response to the 28°C period. In contrast, *eTK* collapsed and *sumo1-1;SUMO2*^*KD*^ showed arrested development after 10 or more days at 28°C. For each combination 8 plants were assessed and the experiment was repeated twice with a similar result. Plants in the white boxes are depicted in (B).

This experiment revealed that development of the vegetative tissue of *sumo1/2*^*KD*^ was stalled as soon as the 28°C-period lasted ten days or more, while growth continued for *sumo1/2*^*KD*^ when this period was only six days (**Fig. 3B, 3C**). After the ten-day warm period, the *sumo1/2*^*KD*^ rosette turned necrotic over the next 20 days while it developed an irregular inflorescence. The warm period of only six days triggered only growth retardation of *sumo1/2*^*KD*^ in comparison to control at 22°C. For *eTK,* a similar observation was made, *i.e.* ten days at 28°C resulted in collapse, while growth of *eTK* resumed after a six-day warm period. Hence, it is not the developmental stage of the mutants, but rather the length of the warm period that defines thermo-lethality for both mutants.

### The shoot apical meristem of *sumo1/2*^*KD*^ becomes highly irregular at 28°C

As *sumo1/2*^*KD*^ and *eTK* failed to resume growth after a warm period, the integrity of their shoot and root meristems was examined using microscopy (**Fig. 4** and **Fig. 5**). Strikingly, integrity of the shoot apical meristem (SAM) and tissue organization were lost when *sumo1/2*^*KD*^ was incubated at 28°C, *i.e.* the cell division patterning and cell size were highly irregular at 28°C, but not 22°C despite the early floral transition (**Fig. 4A-C, 4E,** red arrows). In contrast, the structure of the SAM and cell division patterning were normal for the wildtype (Col-0) and *siz1-2* plants irrespective of the temperature regime given (22°C and 28°C) (**Fig. 4B, 4D**). We also inspected the SAM of *eTK,* but tissue dissection of the *eTK* shoot meristem was only possible till four days at 28°C. Similar to *sumo1/2*^*KD*^, *eTK* had an irregular meristem at 28°C, but not at 22°C (**Fig. 4D**). Patterning of the SAM of *sumo1/2*^*KD*^ was not restored within 7 days upon the return to 22°C. Thus, the SAM is overly sensitive to increased ambient temperatures, and HSFA1 as well as SUMO (conjugation) are critical for this thermo-resilience.

**Fig. 4.**
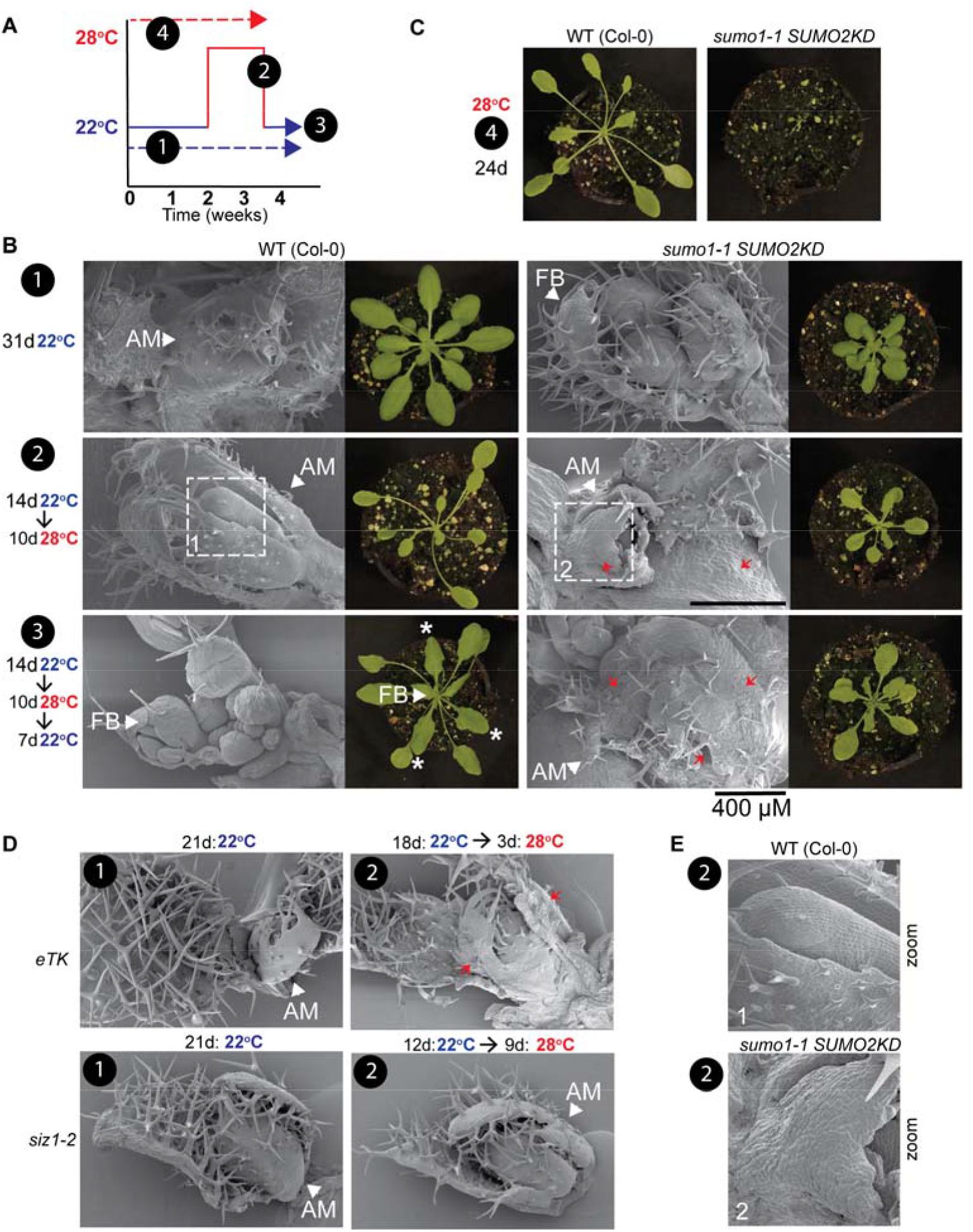
Shoot apical meristem of *sumo1-1;SUMO2*^*KD*^ collapses at 28°C without recovery upon return to 22°C. **A.** Diagram depicting the different temperature regimes (black circles) shown in (B-D). **B.** Cryo-scanning electron microscopy image of the rosette apical meristem (AM) of 24- or 31-day-old plants after exposure to the temperature regimes depicted in (A). The corresponding rosette is displayed on the right. Asterisks (*) marks newly formed leaves upon return to 22°C (without thermomorphogenesis response). FB, floral buds in a bolted rosette. Red arrows highlight the disorganized tissue structure with malformations. For each line and condition at least 8 meristems were inspected with the SEM and the experiment was repeated three times with similar result. WT (Col-0); wildtype background. **C.** Direct germination at 28°C caused seedling lethality of *sumo1-1;SUMO2*^*KD*^ after 24 days, which prevented the SEM analysis. **D.** Same as B, except that the rosette apical meristem of *eTK* and *siz1-2* is shown. The SEM of *eTK* was already inspected after 4 days at 28°C, as after this stage dissecting of the SEM was practically impossible.

**Fig. 5.**
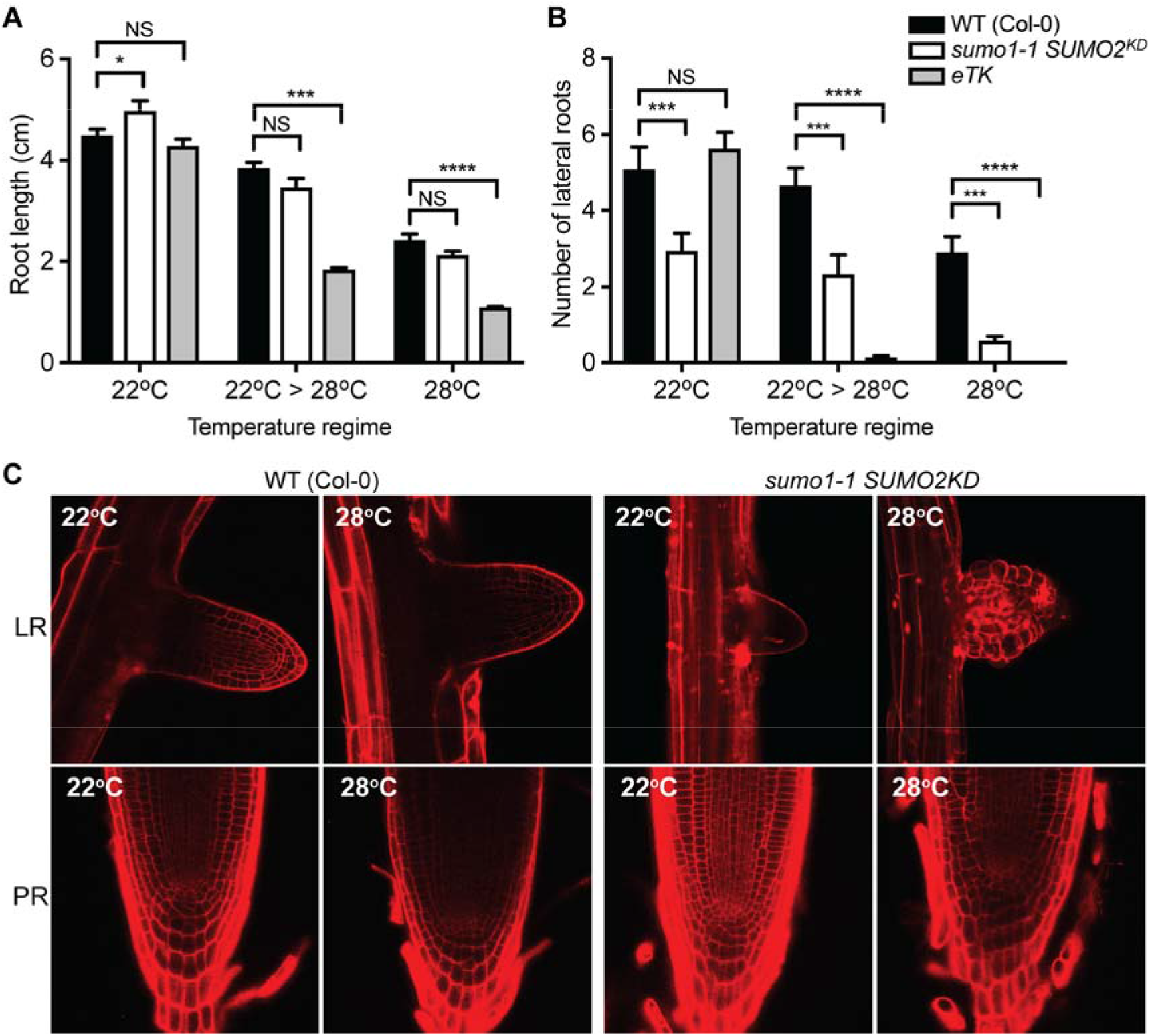
Architecture of the lateral root primordia of *sumo1-1;SUMO2*^*KD*^ is disturbed in response to high ambient temperatures. **A.** Bar graph depicting the average length (±SE) of the primary root of the genotypes Col-0, *sumo1-1;SUMO2*^*KD*^ and *eTK* 12 days post germination at 22°C or 28°C. All plants were germinated and grown for 4 days at 22°C before transferred to new plates for another 8 days at either 22°C, 28°C, or 3 days at 22°C followed by 28°C for 5 days (22C > 28C). In total, the length of approximately 40 roots was measured per line for each temperature regime. Brackets display the result of an ANOVA statistical test followed by Tukey multiple comparison test. Significance results are only shown between the mutants and the wild type control (Col-0) (NS, not significant; *, p<0.05, *** p<0.001, **** p<0.0001). **B.** Similar to (**A),** except that the average number of lateral roots was determined. **C.** Propidium iodide staining showing the architecture of lateral root primordia (LR) and the primary root tip (PR) of 12 to 14-day old seedlings of wildtype (WT) plants (Col-0) and *sumo1-1;SUMO2*^*KD*^ in response to a temperature regime of 22°C or 28°C (experiment was repeated three times with similar results, per condition at least 5 roots were inspected for each experiment).

**Fig. 6.**
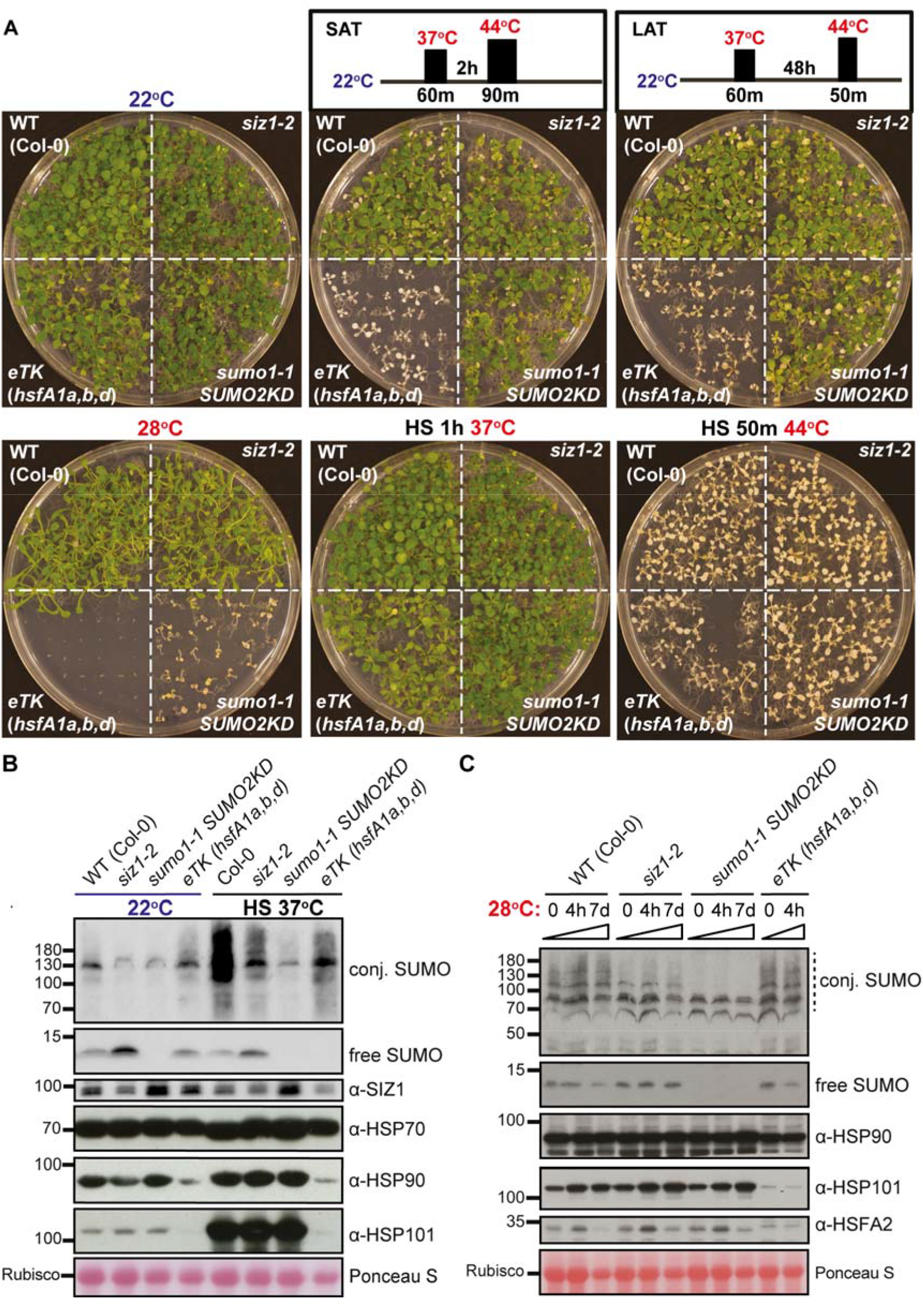
*sumo1-1;SUMO2*^*KD*^ displays a normal acquired heat stress and thermotolerance response. **A.** Heat-sensitive phenotype of wildtype (WT Col-0), *siz1-2, sumo1-1;SUMO2*^*KD*^ and *eTK* (*HsfA1a,b,d*) seedlings in response to different heat shock (HS) regimes (shown at the top). Plants were pre-grown at 22°C for 14 days prior to the treatment indicated. Phenotype of *eTK* is shown as it lacks SAT and LAT. Whereas a single treatment at 44°C for 50 min is sufficient to kill Arabidopsis, WT plants, *siz1* and *sumo1-1;SUMO2*^*KD*^ show normal acclimation when pre-treated at 37°C for 60 min. Experiment was repeated three times with similar result. **B.** Immunoblot showing the conjugated and free SUMO1/2, SIZ1, HSP70, HSP90, and HSP101 protein levels in seedlings (WT Col-0, *siz1-2*, *sumo1-1;SUMO2*^*KD*^ and *eTK*) after 30-min heat stress at 37°C. After a one-hour recovery period, the total protein fraction was extracted. Seedlings were pre-grown for 14 days at 22°C. Ponceau S is shown as a control for equal protein loading. **C.** Immunoblot showing the conjugated/free SUMO1/2 levels, HSP90, HSP101, and HSFA2 levels in seedlings pre-grown at 22°C on plates (0 hrs) and then shifted to 28°C (4 hours or 7 days). Other growth conditions similar to (B). *eTK* samples at 7 days (7d) were not included, as they had collapsed preventing any protein isolation. Ponceau S is shown as a control for equal protein loading.

We also measured whether growth of the main and lateral roots was halted in response to 28°C. Although the main root length of *sumo1/2*^*KD*^ appeared to be shorter at 28°C, this was not significantly different from the growth reduction seen for the wild type control (Col-0) (**Fig. 5A**). This was the case for seedlings that were directly placed at 28°C as well as for plants that experienced first 22°C and then 28°C. This experiment thus reveals that, in contrast to the SAM, growth of the primary root of *sumo1/2*^*KD*^ was not arrested at 28°C. Moreover, *sumo1/2*^*KD*^ developed less lateral roots than the wild type Col-0, but this was independent of the ambient temperature (22°C vs. 28°C) (**Fig. 5B**). In contrast, growth of the primary root of *eTK* was strongly inhibited at 28°C but not at 22°C. Furthermore, lateral root formation was absent for *eTK* at 28°C while normal at 22°C. These data reveal again that SUMO and HSFA1 prevent thermo-lethality, but they likely do so via different mechanisms.

We also inspected development of the root apical meristem in response to high ambient temperatures using confocal microscopy (**Fig. 5C**). Although *sumo1/2*^*KD*^ displayed irregular divisions together with dead cells in the cortex and endodermis (*i.e*. propidium iodide-positive cells), the overall architecture of the root tip appeared to be intact at 22°C and 28°C. This is highly reminiscent of the SUMO E3 ligase mutant *mms21/hpy2* that also shows irregular divisions together with dead cells (Xu *et al.*, 2013). In contrast, the lateral root primordia of *sumo1/2*^*KD*^ were highly irregular at 28°C, but not at 22°C. This deformation of the lateral root primordia of *sumo1/2*^*KD*^ at 28°C resembles the reported lateral root phenotype of the PLETHORA mutant *plt357* (Du and Scheres, 2017; Hofhuis *et al.*, 2013), indicative for a defect in lateral root meristem organization.

### In contrast to *eTK, sumo1/2*^*KD*^ seedlings withstand heat, abiotic and proteotoxic stress

HSFA1 is required for acquired thermotolerance, *i.e. eTK* is highly sensitive to heat stress at 44°C even after pre-exposure to 37°C followed by an acclimation period (Liu *et al.*, 2011). In contrast, SIZ1 is important for (*i*) basal thermotolerance (39°C for up to 4 hours) and (*ii*) the sharp rise in SUMO adduct levels in response to acute heat stress (30 min at 37°C) (Yoo *et al.*, 2006). As *sumo1/2*^*KD*^ mimics to some extent the phenotype of *eTK*, we tested whether acquired thermotolerance is compromised in *sumo1/2*^*KD*^ (**Fig. 6A**). Whereas *eTK* collapsed after a 44°C treatment irrespective of a length of the acclimation period (SAT/LAT), *sumo1/2*^*KD*^ and *siz1* both survived this noxious temperature of 44°C after an acclimation period. This denotes once more that thermo-lethality of *sumo1/2*^*KD*^ at 28°C is not caused by a loss of HSFA1 activity or a failure to regulate the heat-induced transcriptional response (like *HSFA2* upregulation) that promotes heat acclimation. Next, we assessed whether *sumo1/2*^*KD*^ could cope with proteotoxic stress (due to incorporation of L-canavanine for arginine in proteins), water stress (high mannitol) and salt stress, as *siz1* and *eTK* are sensitive to different extents to these conditions (Castro *et al.*, 2015; Liu and Charng, 2013). SUMO adduct levels are also known to increase in response to proteotoxic and abiotic stresses (Conti *et al.*, 2008; Kurepa *et al.*, 2003; Miller *et al.*, 2013). In fact, the notion is that sumoylation is pivotal for the recovery response to nuclear protein damage (at least) in mammals (Liebelt *et al.*, 2019; Seifert *et al.*, 2015). Whereas *eTK* grew poorly on 10 μM L-Canavanine, 300 mM mannitol or 0.75 mM NaCl, both *sumo1/2*^*KD*^ and *siz1* seedlings grew relatively normally (**Supplementary Fig. S2**). We included in this case also *siz1;pad4* and *sumo1/2*^*KD*^;*pad4*, to determine whether the sensitivity would be due to high SA levels, but this did not change the outcome. This experiment thus support the idea that the meristem collapse of *sumo1/2*^*KD*^ at 28°C is not a result of compromised HSFA1 activity.

### *sumo1*/*2*^*KD*^ displays normal HSP protein levels in response to heat stress and warm periods

We also examined the protein levels of known markers of the cellular protein damage response (HSP70, HSP90 and HSP101, and SUMO1/2). To this end, their levels were examined after heat stress (37°C, 30 min). Whereas HSP101 is critical for acquired thermotolerance (Hong and Vierling, 2001), the other proteins have a broad role in the protein damage response. As expected, heat stress triggered a global increase in SUMO conjugate levels in wild type plants, which was largely absent in *sumo1/2*^*KD*^ while strongly reduced in *siz1* and *eTK* (**Fig. 6B**). The latter suggests that *(i)* the HSFA1 protein levels, *(ii)* HSFA1 signaling and/or *(iii)* HSFA1 shuttling to the nucleus is important for the rise in SUMO1/2 conjugate levels in response to heat stress.

The HSP90 and HSP101 levels did not differ between *sumo1/2*^*KD*^*, siz1* and the wild type control (Col-0) after a heat shock. These similar protein profiles indicate again that HSFA1-dependent heat stress response is intact in the SUMO conjugation mutants. In contrast, HSP101 largely fails to accumulate in *eTK* after a heat shock, which matches with the loss of acquired thermotolerance in *eTK*. The levels of HSP90 and HSP101 were also examined in response to a sustained warm period at 28°C (4 hours and 7 days). A warm period had hardly any effect on the HSP90 and HSP101 protein levels in *sumo1/2*^*KD*^*, siz1* and the wild type control **(Fig. 6C**). In contrast, HSP101 levels were again low in *eTK*, while HSP90 levels were slightly reduced. HSFA2 levels displayed a modest transient increase in all lines except for *eTK* after 4 hours at 28°C. This substantiates the other findings that HSFA1 signaling is largely intact *sumo1/2*^*KD*^ and *siz1.* As the wildtype (Col-0) and *eTK* plants did not show a change in their free and conjugated SUMO levels in response to 28°C, we also conclude that a warm period does not lead to a global (persistent) imbalance in SUMO (conjugate) levels.

### The transcriptional response differs between *sumo1/2*^*KD*^ and *eTK* in response to 28°C

Besides that *sumo1/2*^*KD*^ and *siz1* show a delayed and reduced transcriptional response to 28°C linked to a compromised thermomorphogenesis response (Hammoudi *et al.*, 2018), we assessed whether the transcriptional response of *sumo1/2*^*KD*^ to a warm period is mirrored in part by the response of *eTK*. To avoid side effects in the gene expression profiles due to autoimmunity in the two sumoylation-deficient mutants, the *pad4-1* mutation was used again as genetic background. We determined the differentially expressed genes (DEGs) between the lines at each time point in response to the temperature shift (**Fig 7A, Supplementary Dataset S1**). As already reported (Hammoudi *et al.*, 2018), *sumo1/2*^*KD*^*;pad4;* and *siz1;pad4* show a strong overlap in their gene expression profiles 24 and 96 hrs after the temperature shift to 28°C, while the DEGs for *eTK* show less overlap with *sumo1/2*^*KD*^*;pad4* (**Fig 7A**). A principal component analysis (PCA), both on the DEGs and the normalized expression data (*i.e*. without preselection of DEGs) revealed that *eTK* clusters separate from the other three genotypes in this PCA, both at 22 and 28°C (**Fig 7B and Supplementary Fig. S3A**). After 24 h at 28°C, three clusters are visible in the PCA, *i.e*. (1) *pad4,* (2) *eTK,* and (3) *sumo1/2*^*KD*^*;pad4* and *siz1;pad4* combined. At this stage, the first principal component axis, PC1, distinguishes both *eTK* and *pad4* from the sumoylation-deficient mutants. A gene ontology (GO) analysis was then used to identify biological processes that are significantly overrepresented amongst the top 500 DEGs that define the PC axes (**Fig 7C**, **Supplementary Fig. S3B, Supplementary Dataset S2-S4**). For PC1 we find an overrepresentation of the GO terms ‘cell division’, ‘DNA replication’, ‘chromosome organization’, ‘(mitotic) cell cycle’, ‘microtubule movement’ and ‘DNA metabolic process’ of the PCA. These GO terms were also overrepresented at the late time point (96 h at 28°C), but they now contribute to PC2, which separates *siz1;pad4* from *pad4-1* while *sumo1/2*^*KD*^*;pad4* takes an intermediate position. These GO terms were not significantly overrepresented for PC1 and PC2 prior to the temperature shift (22°C). This suggests that the loss of SIZ1-dependent sumoylation alters the transcriptional response linked to plant growth when the temperature increases. This is in line with the fact that the thermomorphogenesis response is compromised in *siz1-2.* When looking at the other axis (PC1 at T1-24h and PC2 at T2-96h), we see that the GO terms ‘response to stress’, ‘response to abiotic stimulus’ ‘defense response’, ‘innate immune response’, ‘response to temperature are overrepresented. These results did not change when we performed an unbiased PCA on the top 500 genes that contribute to the PC loadings based on the entire gene chip dataset (**Supplementary Fig. S3B** and **Supplementary Dataset S4**).The majority of the DEGs was detected for *eTK* after 96 h at 28°C when this mutant already collapses (visualized by PC1 at this stage that explains 73/78%). Importantly, none of the GO terms identified for PC1 at T=1 or PC2 for T=4 (24 h and 96 after the shift 28°C, respectively) was already significant prior to shift 28°C. Evidently, the transcriptional responses of *sumo1/2*^*KD*^ and *eTK* differ despite the fact that both are hypersensitive for a prolonged period at 28°C.

**Fig 7.**
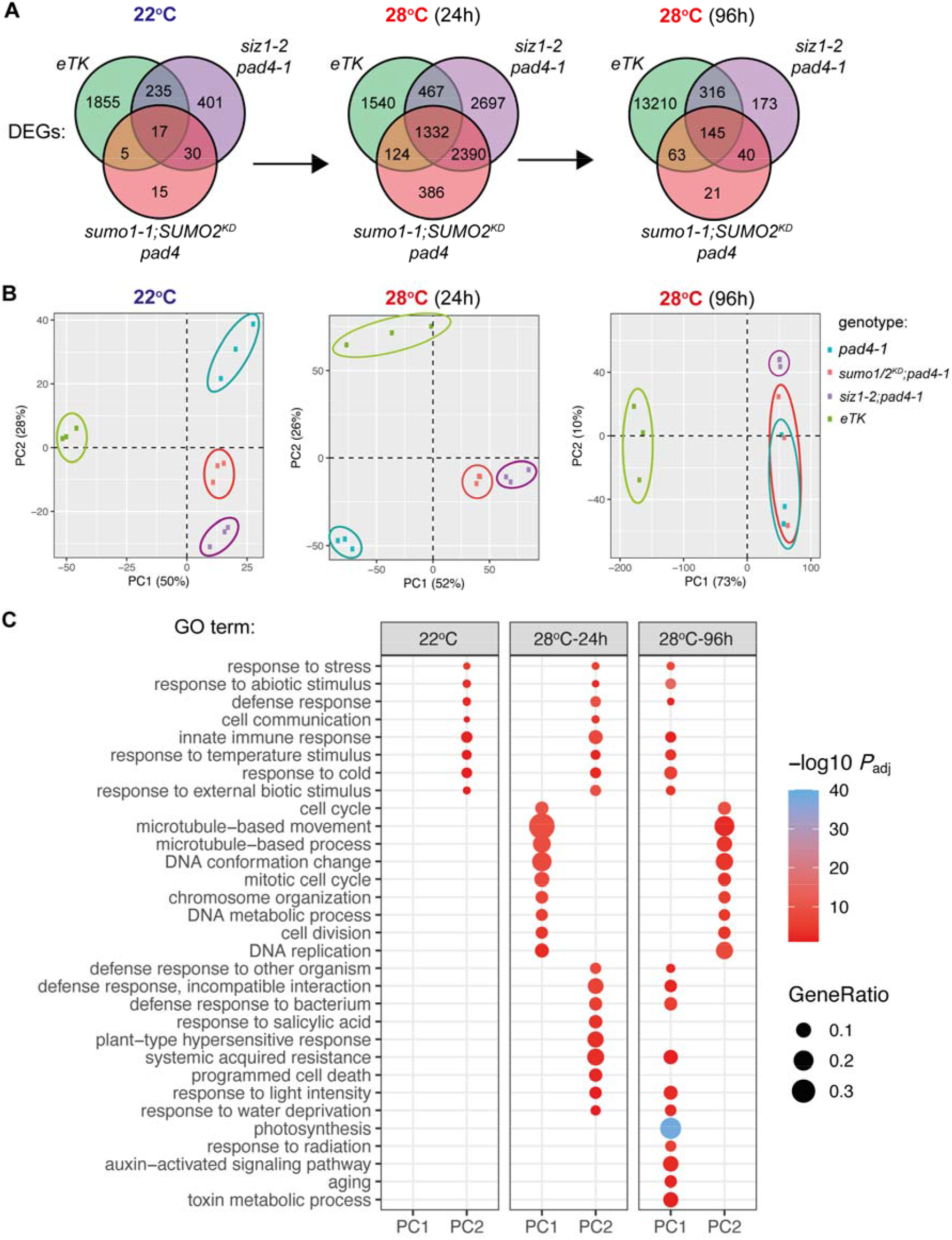
Transcriptional response of *sumo1-1;SUMO2*^*KD*^ and *eTK* differs in response to a sustained warm period at 28 degrees Celsius. **A.** Venn diagrams showing the total number of differentially expressed genes (DEGs) detected at the different time points for each genotype (compared to *pad4-1*) and their overlap. The overlap in the transcriptional response after 24 hours at 28°C between *sumo1/2*^*KD*^*;pad4* and *siz1;pad4* is largely due to a delayed thermomorphogenesis response in both plant lines (Hammoudi *et al.*, 2018). All DEGs passed a FDR of *q*-value < 0.01. **B.** Principal component analysis (PCA) of DEGs detected showing that *eTK* responds distinct from both SUMO-deficient mutants (*sumo1/2*^*KD*^*;pad4* and *siz1;pad4* and the control (*pad4-1*) (n=3 for each genotype/time point). **C.** Dot plot depicting enriched gene ontology (GO) terms for the top 500 DEGs (either overexpressed or downregulated) that contribute to the loading of the principal component (PC) axes shown in panel B. Dot size indicates the *k*/*n* ratio (“gene ratio”), where *k* is the number of genes participating in this GO term, and *n* is the total number of genes annotated for this GO term in the genome. Dot color indicates the adjusted p-value of the enrichment test (hypergeometry test with Yekutieli FDR correction, *P*adj <0.01). GO terms shown were manually selected to best represent the biological processes impacted for interdependent GO-terms.

## DISCUSSION

Here we report that SUMO1 and −2 combined are essential for Arabidopsis to endure sustained warm periods of 28°C (as depicted in a model in **Supplementary Fig S4**). None of the other components of the SUMO (de)conjugation machinery tested appeared to be required for thermo-resilience—alone or in combination. We also found that poly-SUMO chain formation is not necessary to sustain warm temperatures. This implies that SUMO thermo-resilience apparently depends on mono-sumoylation of one or more substrates. At the tissue level, we found that in particular the integrity of the shoot apical meristem and lateral root primordia was lost in the SUMO1/2 (conjugation) knockdown mutant when plants were grown at 28°C. This deformation of the lateral root primordia of *sumo1/2*^*KD*^ resembled the lateral root phenotype of the PLETHORA mutant *plt357* (Du and Scheres, 2017; Hofhuis *et al.*, 2013). These PLETHORA transcription factors are required for the formative periclinal cell divisions that initiate lateral root primordia. Apparently, SUMO has a role in lateral root initiation. In the *sumo1/2*^*KD*^, this process is foremost disturbed at increased temperatures, possibly via a pathway that involves PLT3/PLT5/PLT7, which together also control Arabidopsis phyllotaxis (Hofhuis *et al.*, 2013). The role of SUMO1/2 for meristem integrity at elevated temperatures was independent of EDS1/PAD4 and accumulation of the defense hormone salicylic acid by SID2. Thus, thermo-resilience is a third aspect of the pleiotropic phenotype of the *sumo1/2*^*KD*^ mutant—besides inhibition of SNC1-dependent autoimmunity and thermomorphogenesis (Hammoudi *et al.*, 2018; van den Burg *et al.*, 2010).

Our finding that SUMO confers thermo-resilience to warm periods was previously reported for HSFA1 (Liu *et al.*, 2011). In general, HSFA1 is regarded to be the master regulator of the heat-stress response. Nonetheless, we see differences between *sumo1/2*^*KD*^ and *eTK*, suggesting that the two proteins act in different (regulatory) processes. First, SUMO was not required for both short- and long-term acquire thermotolerance while HSFA1 is. Second, accumulation of the chaperones HSP90 and HSP101 but also the transcription factor HSFA2 was largely intact in *sumo1/2*^*KD*^ and *siz1* in response to heat stress or increased ambient temperatures, but not in *eTK*. Both HSP101 and HSFA2 protein levels are well known markers for heat-stress induced nuclear activity of HSFA1 (Busch *et al.*, 2005; Liu *et al.*, 2011). This experiment thus suggests that HSFA1 is still responsive when heat stress (protein damage) is applied to *sumo1/2*^*KD*^ and *siz1.* The transcriptional profile of *sumo1/2*^*KD*^*;pad4* resembles also more that of *siz1;pad4* than *eTK* in response to warm conditions. We also noted that *eTK* does not germinate at 28°C, while *sumo1/2*^*KD*^ does germinate but it collapse within 2 weeks post germination. Combined these findings argue that the HSFA1 regulatory pathway is largely intact and responsive in the sumoylation-deficient mutants *sumo1/2*^*KD*^ and *siz1*.

Mammals express two close homologues of Arabidopsis HSFA1, called HSF1 and HSF2. HSF1 and −2 undergo stress-induced sumoylation, which modulates their transcriptional activity and DNA affinity (Akerfelt *et al.*, 2010; Anckar *et al.*, 2006; Goodson *et al.*, 2001; Hietakangas *et al.*, 2003; Hong and Vierling, 2001). Also in plants, different HSF family members (HSFA1b, HSFA1d, HSFA2 and HSFB2b) and some downstream targets (*e.g*. DREB2A) are subject to sumoylation (Augustine and Vierstra, 2018; Cohen-Peer *et al.*, 2010; Miller *et al.*, 2010; Rytz *et al.*, 2018; Wang *et al.*, 2020). Of note, none of these HSF transcription factors appeared to be sumoylated in a SIZ1-dependent manner (Rytz *et al.*, 2018). The latter would favor that SCE1 directly targets these HSFs for SUMOylation. Yet the biological consequence of HSFA1 sumoylation remains unknown. In the case of tomato HSFA2, sumoylation was suggested to negatively control its transcriptional activity (Cohen-Peer *et al.*, 2010). As the Arabidopsis *hsfa2* mutant showed normal thermo-resilience to 28°C, it is unlikely that HSFA2 is important for the here seen SUMO-mediated thermo-resilience. Also many chaperones including HSP90 are SUMO modified in mammals and yeast (Panse *et al.*, 2004; Pountney *et al.*, 2008; Zhou *et al.*, 2004). In Arabidopsis HSP90 was shown to inhibit HSFA1 by sequestrating the protein outside of the nucleus (Yoshida *et al.*, 2011; Zou *et al.*, 1998). Hence, HSP90 directly acts on HSFA1 in a negative feedback loop, although in absence of HSP90, other factors may be required to strongly activate HSFA1 in the nucleus during stress conditions (Yoshida *et al.*, 2011). We observed that the rapid increase in SUMO conjugates due to acute heat stress in part also depends on HSFA1. Possibly, HSFA1 sequesters SUMO and/or the E2 enzyme under normal conditions. Alternatively, activation and nuclear translocation of HSFA1 might impact the SUMO conjugate levels by promoting SUMO conjugation and/or by reducing SUMO protease activity. Clearly, future studies should determine how the HSFA1/HSP90 regulatory network affects SUMO and how SUMO affects the plant proteostasis in response to acute heat stress, but also a mild increase of the ambient temperature. At this stage we cannot rule out that SUMO conjugation modulates HSFA1 activity directly or part of its downstream responses. Clear follow up questions are: does SUMO modification of HSFA1 impact the interaction with HSP90 in the cytosol or the nuclear functions of HSFA1 when bound to *HRE* cis-regulatory elements? And why is the meristem sensitive to loss of HSFA1 and SUMO?

## Abbreviations

DEG: differentially expressed gene
GO: gene ontology
HSF: Heat Shock transcription Factors
HSP: Heat Shock Protein
LAT/SAT: long/short-term acquired thermotolerance
LD_50_: median lethal dose
PCA: principal component analysis
ROS: reactive oxygen species
SAM: Shoot apical meristem
SUMO: Small ubiquitin-like modifier
sumoylation: SUMO attachment to substrates.

## Supplementary data

Supplementary data are available at *JXB online*.

*Table S1.* Resources used in this study

*Table S2.* Primers used in this study

*Table S3.* Details for Arabidopsis plant genotyping

*Fig. S1.* Growth phenotype of different heat-sensitive mutants at 22°C and 28°C.

*Fig. S2.* Response of *sumo1/2*^*KD*^ to different abiotic and proteotoxic stresses.

*Fig. S3.* Unbiased gene ontology enrichment analysis of the gene expression data in response to warm ambient temperatures for *sumo1/2*^*KD*^*;pad4-1*, *siz1-2;pad4* and triple *HSFA1* mutant *eTK* in comparison to *pad4-1*.

*Fig. S4.* Diagram depicting the role of SUMO and HSFA1 in thermo-resilience and heat stress.

*Dataset S1.* Datasheet with the DEGs detected in *sumo1/2*^*KD*^*pad4, siz1 pad4,* and *eTK* in comparison to the control plant (*pad4-1*) at the different time points (22°C, 28°C-24h and 28°C-96h).

*Dataset S2.* Datasheet with the top500 DEGs that contributed to loading of PC1 and PC2 at each time points (22°C, 28°C-24h and 28°C-96h); Gene ID list was used to perform the GO term analysis shown in **Fig 7C**.

*Dataset S3.* Datasheet with the top500 genes that contributed to loading of PC1 and PC2 at each time points (22°C, 28°C-24h and 28°C-96h); Gene ID list was used to perform the GO term analysis shown in **Fig S4B**.

*Dataset S4.* Datasheet with the selected Gene ontology terms for the (a) top500 genes for the DEGs (**Fig 7C**) and the top500 genes unbiased that contributed to the loading of the PC axis (**Fig S4B**)

## Acknowledgments

We thank Like Fokkens, Jeffrey Ham, Ludek Tikovsky, Harold Lemereis for help with gene expression analyses, genotyping, and plant maintenance, respectively. We express our gratitude to all colleagues who supplied seeds and materials. This work was supported by grants of Netherlands Scientific Organisation (ALW-VIDI 864.10.004 to HvdB; TTW14948 to TH). We thank the Van Leeuwenhoek Centre for Advanced Microscopy (SILS-University of Amsterdam) for use of microscopy facilities.

## Author contributions

HB and VH designed the research with input from all authors.

VH, BB, and HB performed and analyzed plant growth experiments.

MK, BB and HB performed and analyzed the root growth experiments.

HB, MG, TH and BB performed the SEM experiments.

BB performed the immunoblot analysis and abiotics stress plate assay

MJ and HB performed the gene expression analysis. BB prepared the RNA samples.

BB prepared and analyzed the protein data.

HB wrote the final manuscript with input from all authors

## Data availability

All data supporting the findings of this study are available within the paper and its supplementary materials published on line. The original microarray data that support the findings of this study are openly available in the GEO database at NCBI (https://www.ncbi.nlm.nih.gov/geo) under accession number GSE97641.

